# Conserved odor detection and divergent behavioral valence in ecologically distinct drosophilid flies

**DOI:** 10.64898/2026.05.22.727105

**Authors:** Venkatesh Pal Mahadevan

## Abstract

Competition among living organisms often drives selection for the exploitation of new ecological niches. Successfully colonizing these niches requires the evolution of specialized physiological adaptations. In drosophilid flies, which depend heavily on their sense of smell, such adaptations frequently involve the modification of specific olfactory circuits to detect and respond to niche-defining odorants. Although evolutionary patterns within the *Sophophora* subgenus have been studied to a great extent, the peripheral circuit evolution, as well as the extent to which the peripheral code is conserved in other subgenera, remains poorly understood. Here, I compared the antennal basiconic olfactory coding patterns in *D. melanogaster, D. virilis* and *D. busckii*, belonging to the subgenera *Sophophora, Drosophila* and *Dorsilopha*, respectively. Despite their substantial evolutionary divergence (∼40 Mya), I found partial conservation of basiconic peripheral coding, together with several strongly conserved ligand responses across species. I then tested whether conserved peripheral detection also corresponds to conserved behavioral valence. These experiments revealed that species can exhibit divergent behavioral responses to the same ligand, even when peripheral detection is conserved. Finally, I describe four individual cases of circuit evolution across selected species pairs, providing independent case studies of peripheral olfactory evolution.

## Introduction

Animals that rely on their sense of smell must efficiently perceive and process the complex olfactory landscape of their surroundings. Species of the genus *Drosophila*, in particular, have been studied extensively in this context, as they depend heavily on olfaction and occupy a wide range of ecological niches [1]. The differences between the niches are often reflected and paralleled by corresponding olfactory adaptations [2–8]. In addition to their general dependence on the sense of smell, the genetic information for these species is widely available [9–11]. This combination of genetic accessibility, along with diverse niche occupations, makes the *Drosophila* genus a powerful system to study olfactory evolution. As one can expect, the olfactomes of such ecological niches differ significantly from each other and prompt investigations of how the olfactory system has evolved in order to detect and process such niche-specific odorant blends.

In *Drosophila*, evolutionary shifts in the olfactory system can be studied effectively at the level of peripheral odor detection where ligands are selectively detected by the odorant receptors (ORs) expressed on olfactory sensory neurons (OSNs) in the fly antenna. The combined responses of OSNs to one or more ligands form the peripheral olfactory code. Evolution of such peripheral olfactory code can be investigated by studying changes at the molecular (OR) or functional (OSN) level. Several studies have examined the molecular evolution of the complete OR repertoire across multiple drosophilids, revealing interesting trends in OR gene duplication as well as loss-gain events [9–11]. Comparative databases generated from these studies have further enabled investigations of changes in local, individual olfactory circuits. For example, ORs such as Or22a, Or85a and Or67a in *Dmel* as well as in its close relatives, including *D. sechellia* and *D. suzukii* have been identified as hotspots for rapid evolutionary changes, where shifts in their tuning properties link to differential behaviors and host specialization [12–15]. Taken together, studies investigating molecular and circuit evolution highlight the evolutionary contribution of individual olfactory circuits in niche-specific adaptations [16].

Ecological niches occupied by drosophilids are often rich in chemically diverse compounds, making it essential to understand whether and how peripheral coding of these complex odor bouquets evolves across species. A few studies have investigated the evolution of the peripheral olfactory code across closely related species [14,17–20]. Surprisingly, it seems that the peripheral code is to a great extent conserved across closely related species, where minor shifts in selected receptors/circuits give rise to novel behaviors. For example, the global peripheral code within subgenus *Sophophora* is largely conserved [17–19,21] yet changes pertaining to Or22a and its downstream circuit explain host specialization of *Sophophora* subgroup species [2,4,6,12,22]. Although these studies have greatly advanced our understanding of the functional evolution of the fly olfactory system, they share a major limitation: all the species examined are associated with fruit hosts of various types and ripening stages, thereby excluding volatiles from a range of non-fruit ecological niches. In summary, the evolution of peripheral olfactory coding across species occupying diverse ecological niches and across different subgenera remains poorly understood.

I have recently expanded our portfolio by studying the olfactory neuroecology of *D. virilis* (*Dvir*) [8] and *D. busckii* (*Dbus*) [7], belonging to the subgenera *Drosophila* and *Dorsilopha*, respectively. These species have unusual niche preferences as they breed from rotting tree sap [23,24] and rotting vegetables [25], respectively. In addition to demonstrating some key niche-specific odorant preference events, I have also established the peripheral olfactory code in these two species by screening basiconic sensilla on the antenna. Given that *Dbus* and *Dvir* are evolutionarily distant from each other and from *Dmel*, I sought to identify ligands that maintain their behavioral valence across these divergent species. First, I compared the antennal basiconic olfactory codes among the aforementioned species. I identified multiple conserved responses to several key odorants. However, the behavioral valence of these conserved ligands varied across species, which could be partially explained by their distinct ecological backgrounds. Finally, I present specific functional examples, including loss of response, redundant response, altered sensitivity to key odorants, and reshuffling of OSN responses within a given sensillum type, as independent and striking case studies across the three species. Taken together, this study provides new insights into olfactory coding patterns in drosophilid flies that occupy non-overlapping ecological niches.

## Results

### Peripheral olfactory coding evolution across three subgenera

I studied *D. melanogaster* (*Dmel*), *D. virilis* (*Dvir*), and *D. busckii* (*Dbus*) as representatives of the subgenera *Sophophora, Drosophila*, and *Dorsilopha*, respectively (Fig. 1a). These three species, which diverged approximately 40 Mya, occupy distinct ecological niches: fermenting fruits, slime flux, and rotting vegetables, respectively (Fig. 1a). To characterize these niches, I leveraged chemical profile data from multiple ecologically relevant substrates, including several fermenting fruits, rotting vegetables, and naturally collected slime flux samples reported in previous publications [7,8]. I then performed a principal component analysis (PCA) on the raw chromatograph trace using XCMS Online [26] to establish the qualitative differences in the chemical odor space of the three substrate types. The PCA revealed distinct, non-overlapping distributions of fruit, vegetable, and slime flux-derived odors, indicating a clear separation of niches based on their odor profiles (Fig. 1b).

**Figure 1:**
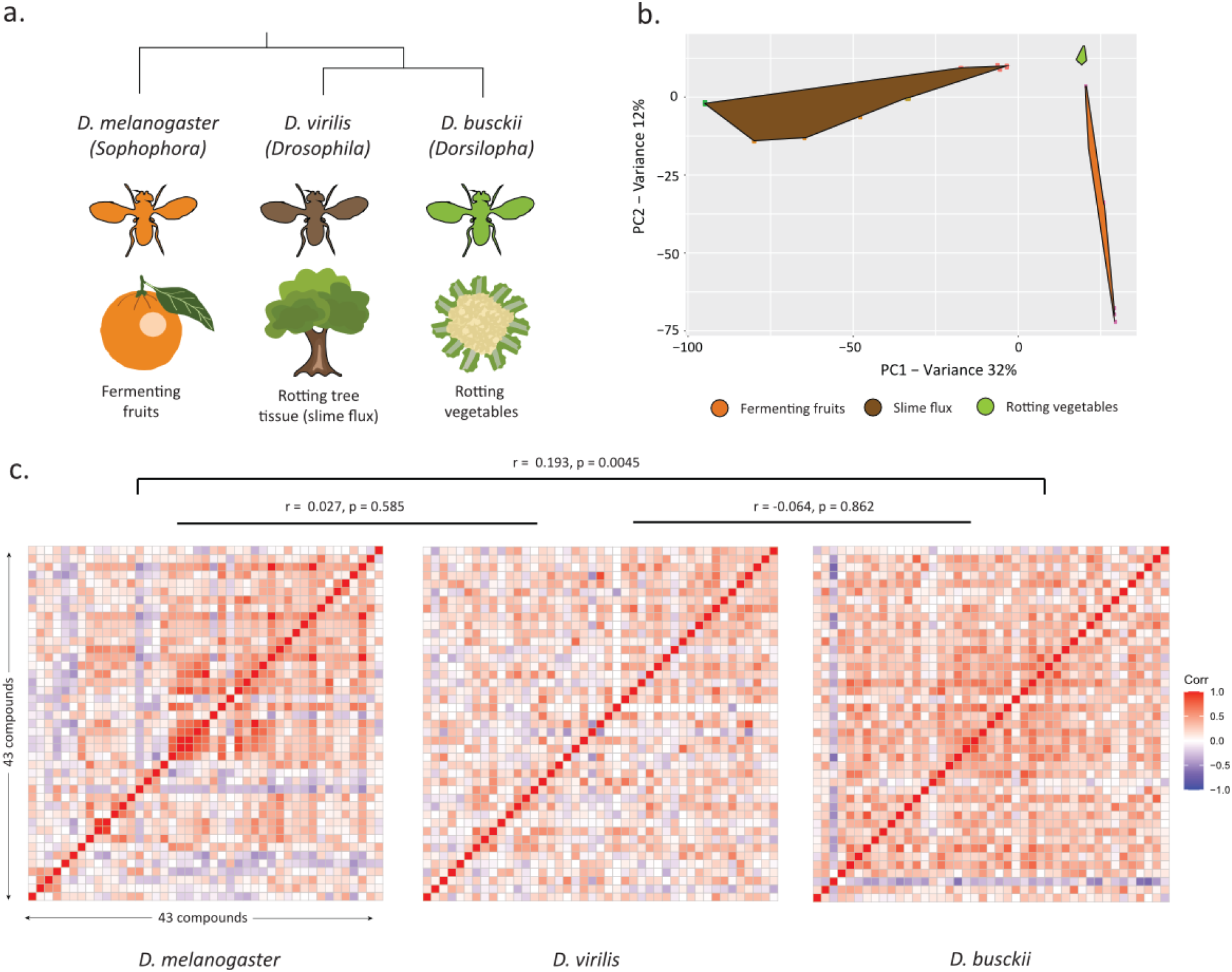
a. Cartoon representation of the three species tested in this study, along with their respective ecological niches. Phylogeny only for representation and does not correspond to the evolutionary timescale. b. Principal component analysis (PCA) of chromatograms of all odor spaces collected from the ecological niches corresponding to the three species tested. c. Spearman correlation matrices were generated for each species, where each block represents a pairwise correlation between SSR responses for odors coded by the different functional neuron types. The significance of the correlation matrices was tested using the Mantel test, and the values are mentioned on top.

### Comparative coding of odor spaces across species

I hypothesized that the distinct odor spaces identified for fruits, vegetables, and slime flux must also be coded in a significantly different manner by OSN types across the three species. To test this, I leveraged single sensillum recording (SSR) datasets (supplementary table 1) from previous studies in which a panel of 43 chemically diverse compounds (encompassing volatiles from all three ecological habitats) was used to screen OSNs present in antennal basiconic sensilla across *Dmel, Dvir* and *Dbus* [7,8]. First, I examined whether the global coding pattern of odorants by basiconic sensilla varied between species. For each species, correlation indices were calculated between all pairs of responses to the compounds in the screening panel (Fig. 1c). Surprisingly, correlation matrices generated by comparing odor pairs revealed modest but significant similarity between *Dbus* and *Dmel* coding patterns (Mantel r = 0.193, Holm-adjusted p = 0.0045; bootstrap 95% CI: 0.122–0.367; n = 43 odors). In contrast, comparisons involving *Dvir* were not significant, wherein the comparisons between *Dbus* vs *Dvir* showed no similarity (Mantel r = −0.064, Holm-adjusted p = 0.862; 95% CI: −0.103–0.135), while *Dmel* vs *Dvir* was also not significant (Mantel r = 0.027, Holm-adjusted p = 0.585; 95% CI: −0.020–0.222). These results suggest partial conservation of population-level peripheral coding structure between *Dbus* and *Dmel*, divergence in comparisons involving *Dvir*.

### Nomenclature

In the combined dataset (Supplementary table 1), I have prefixed each sensillum type with the originating species: M, V, and B for *Dmel, Dvir* and *Dbus*, respectively. Sensillum types were numbered independently for each species in the order in which they were first identified during screening experiments. Thus, for example, Vab7 and Bab7 represent distinct sensillum types and should not be assumed to be orthologous.

### Conservation of coding of key ligands

I next performed hierarchical clustering of OSN responses across all three species (Fig. 2). This analysis revealed that while some odorants are perceived in a highly conserved fashion (highlighted in red, Fig. 2), most OSN types did not follow a stereotypical clustering pattern. In addition, a subset of OSNs showed only weak responses to a limited number of odors and did not respond strongly to any compound in the panel, rendering them “orphan” OSNs without a strong, unique diagnostic odor identity. It is noteworthy that I matched OSN classes across species based on shared diagnostic responses, as molecular receptor identification in non-model species remains technically challenging. However, functional matching does not necessarily indicate expression of orthologous receptors. Clustering of two or more OSN classes could, in principle, also reflect comparisons of non-orthologous OSNs converging on a similar response profile.

**Figure 2:**
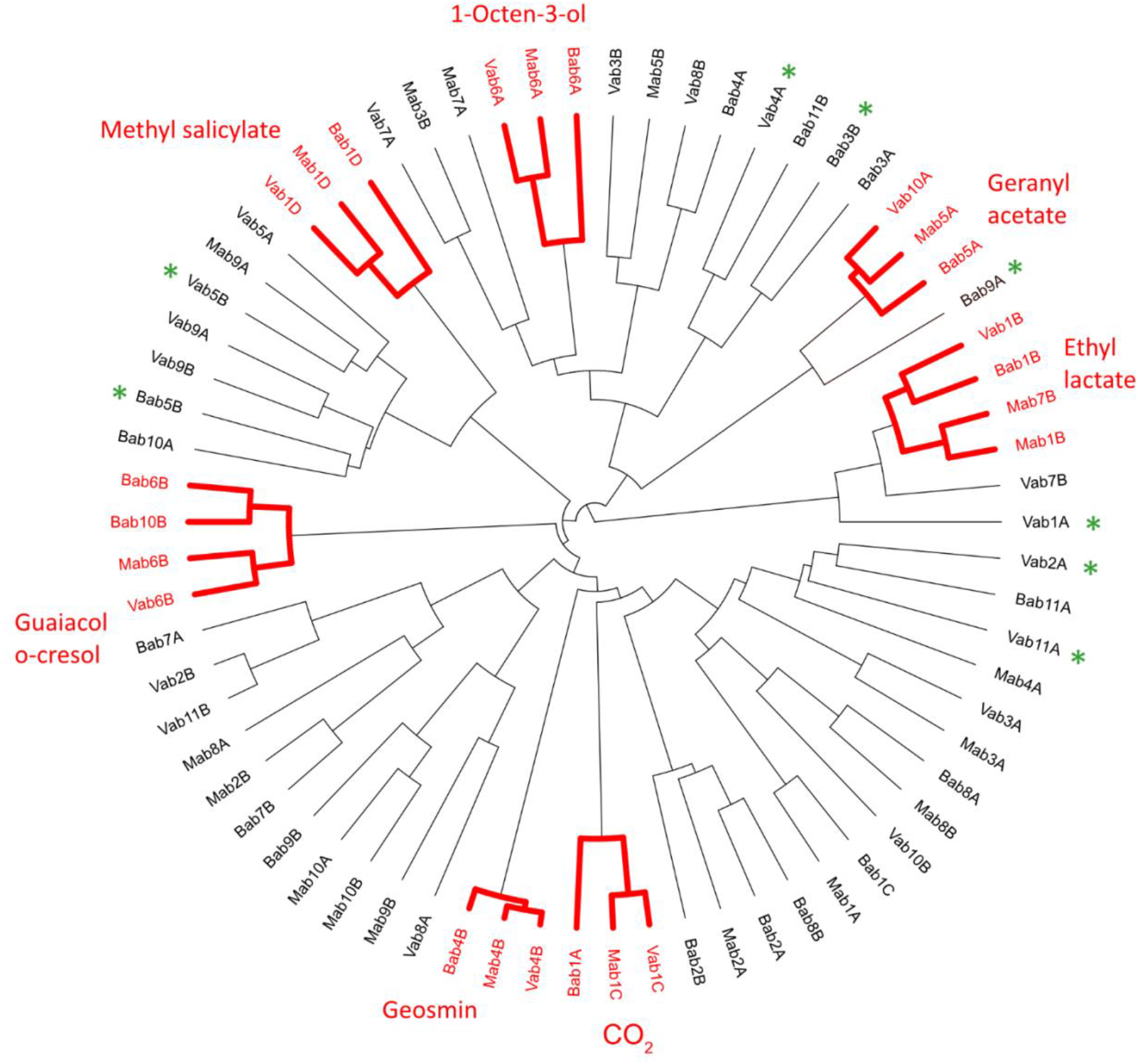
Hierarchical clustering dendrogram. Each abbreviation denotes a sensillum type based on its response to a panel of 43 diverse chemical stimuli. The prefixes M, B and V denote the three species, *Dmel, Dbus* and *Dvir* respectively. The functional types that are highly conserved across the three species are highlighted in red. Functional types that had distinct responses without any strong diagnostic odor are identified by green asterisks and do not have a strong diagnostic ligand.

Interestingly, ligands that were conserved across subgenera (e.g., guaiacol, geosmin, CO_2_, geranyl acetate, methyl salicylate) were primarily encoded by narrowly tuned OSNs in *Dmel* [27,28]. This observation prompted me to ask whether such conservation at the level of detection is also reflected in behavior. To test behavioral conservation, I performed dual-choice preference assays with each of the conserved ligands across the three species, using ecologically relevant concentrations (10^−4^ v/v). Each odorant elicited species-specific valence responses (Fig. 3a). For example, guaiacol was highly attractive to *Dvir* but neutral to both *Dmel* and *Dbus*. Conversely, geosmin, a harmful microbial volatile [27], was aversive to *Dmel* and *Dvir*, but surprisingly did not repel *Dbus*. I then investigated whether these behavioral differences reflected differences in OSN firing strength. However, direct comparisons of functionally matched OSNs across the three species revealed highly similar firing rates (Fig. 3b), although the molecular orthology of these OSNs remains to be confirmed.

**Figure 3:**
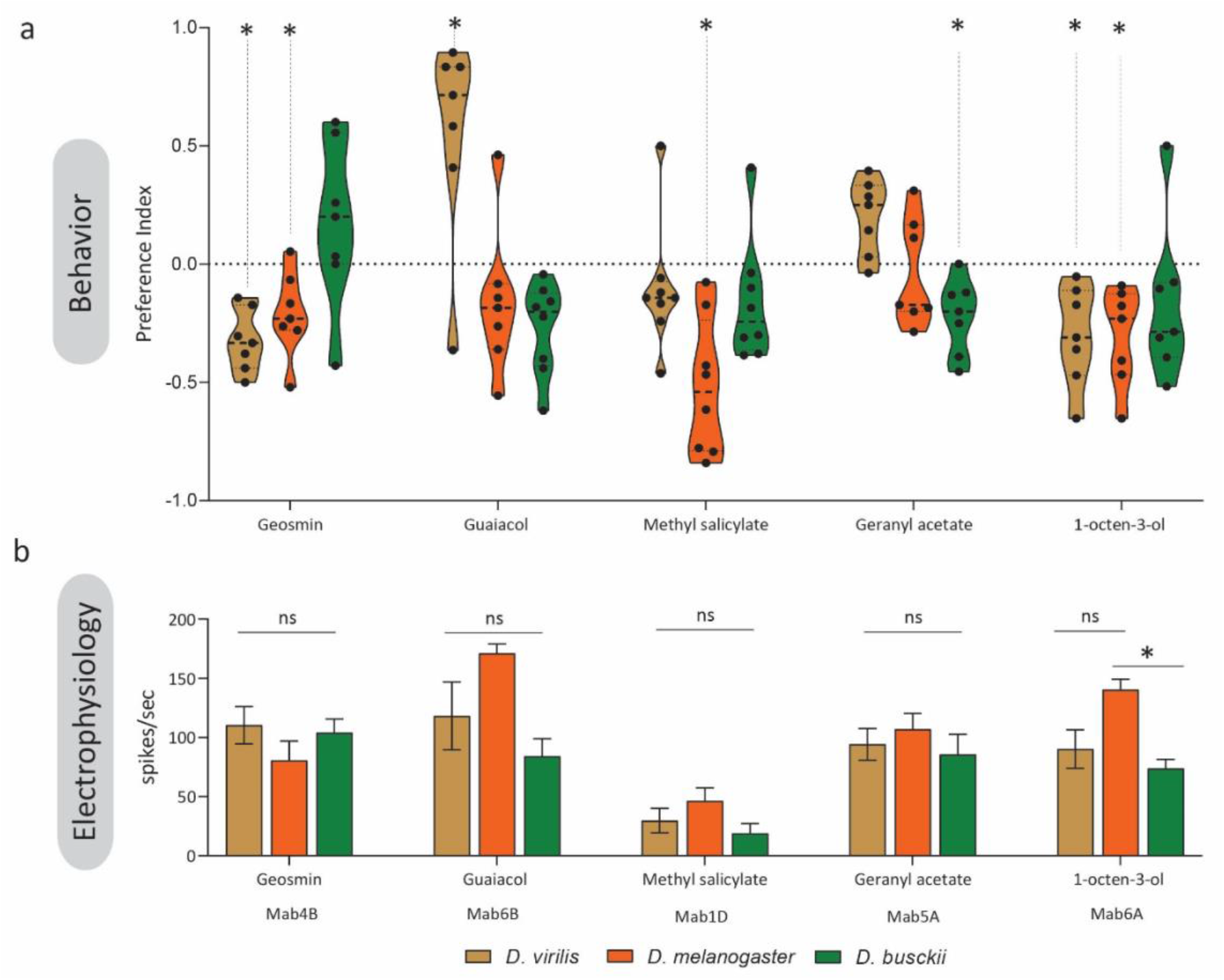
Behavioral and electrophysiological responses to conserved ligands. a. Preference indices representing the choice of each species when tested between individual compounds (10^-4^ v/v) and control (mineral oil). Significance was calculated between the preference index and neutrality (0). * p < 0.05. b. Response strengths of individual OSN types to 10^-4^ v/v concentration of individual compounds. The comparison was carried out between OSN types known in *Dmel*, along with the functionally matched OSN types identified in this study (see Fig. 2). n = 3-5

### Case studies of evolutionary shifts

Beyond the broadly conserved ligands, I identified several examples of species-specific shifts that could be grouped into four categories: (a) shifts in OSN response dynamics, (b) changes in OSN sensitivity, (c) redundant responses, and (d) loss of response.

#### a. Shift in OSN response dynamics

In *Dbus*, the Bab1 sensillum (comparable to Mab1 and Vab1 in *Dmel* and *Dvir*) contains four OSNs and shares diagnostic responses to ethyl lactate, methyl salicylate, and CO_2_ (Supplementary tables 1). However, in Bab1, the large-amplitude Bab1A OSN responds to CO_2_, whereas in *Dmel* the shorter-amplitude Mab1C OSN responds to CO_2_ (Fig. 4a). Likewise, ethyl lactate was detected by Bab1C in *Dbus* but by Mab1B in *Dmel*. Responses to methyl salicylate were conserved in the smallest neuron (D cell) in both species, albeit weaker in *Dbus*.

**Figure 4:**
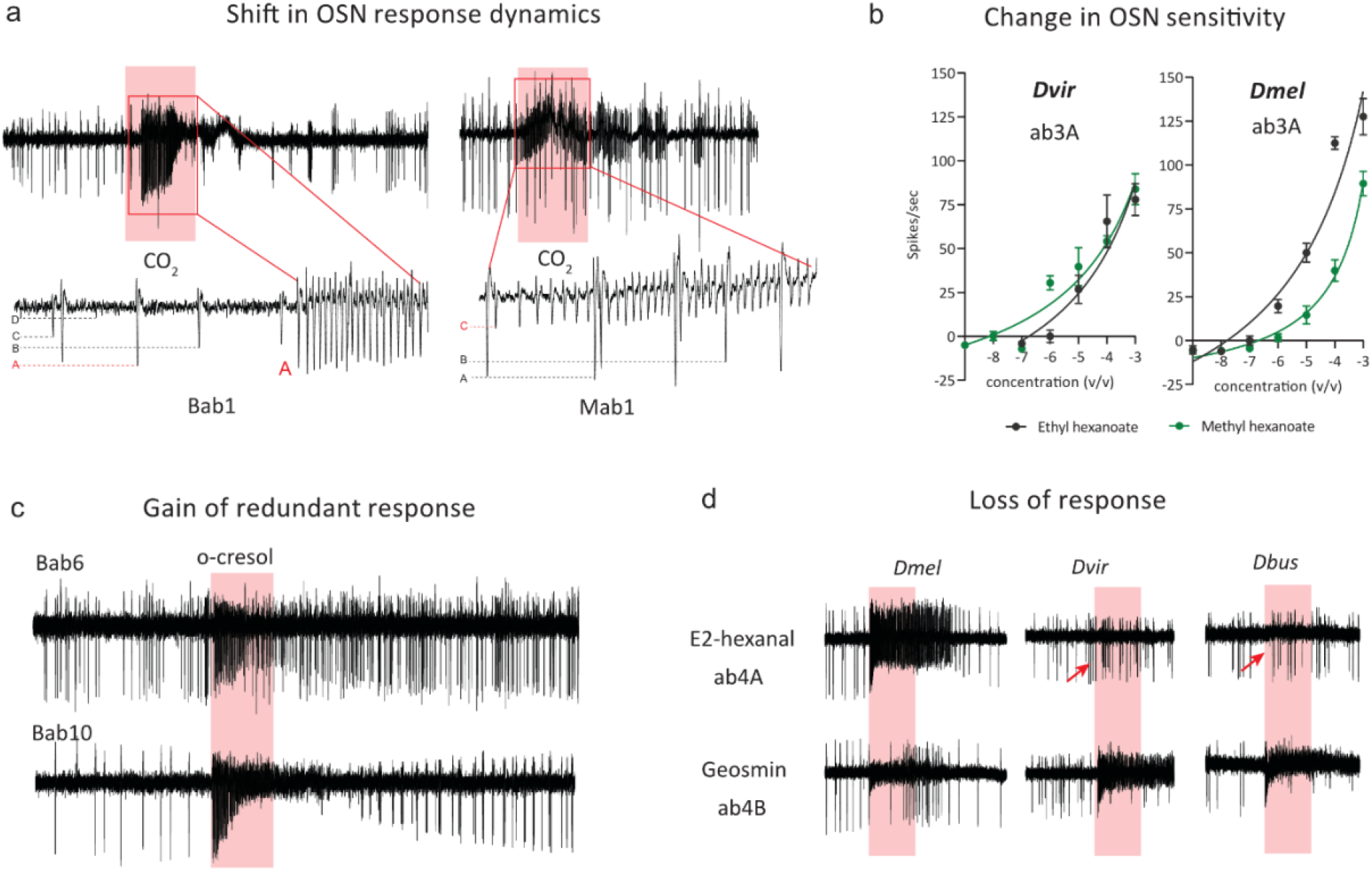
Individual case studies of directional olfactory evolution. a. (left) Sample trace from the Bab1 OSN type, where responses from 4 OSN types can be distinguished based on their relative amplitudes. The largest A OSN responded to CO_2_. On the right, a sample trace from an Mab1 OSN type, where the smaller C neuron is activated by CO_2_ stimulation. Lower panels show a zoomed-in view of individual spike amplitudes in both examples. b. Dose-response experiments involving testing of sensitivity towards methyl and ethyl hexanoate esters in *Dvir* and *Dmel* from functionally matched OSN types (Vb3A vs Mab3A). c. Representative traces of two *Dbus* OSN types (Bab6 and Bab10) responding strongly to o-cresol. d. the lower panel shows representative traces of functionally matched sensillum types based on the common diagnostic odor, geosmin (Mab4B, Vab4B and Bab4B) across the three species. The neighbouring OSN type in the same sensillum shows variation in the diagnostic odor response. In the upper panel, the Mab4A OSN type responds strongly to E2-hexanal, while this response is absent in the case of Vab4A and Bab4A (red arrowheads).

#### b. Change in OSN sensitivity

The Vab3 and Mab3 sensilla of *Dvir* and *Dmel* each contained an OSN (the larger spike amplitude A OSN) tuned to esters such as ethyl hexanoate and methyl hexanoate (supplementary table 1). Comparative responses indicated a reversal of sensitivity between the two species. Dose-response experiments demonstrated that Vab3A responded more strongly to methyl hexanoate, whereas Mab3A responded preferentially to ethyl hexanoate (Fig. 4b). Notably, Mab3A OSNs were more sensitive overall than Vab3A, and no Mab3A-like sensillum (responding to esters) was identified in *Dbus*.

#### c. Redundant response

OSN type ab6B showed conserved responses to o-cresol (also known as 2-methyl phenol) across all three species, clustering together due to strong responses to guaiacol and o-cresol (Fig. 2). However, in *Dbus*, a second sensillum type (Bab10) was identified in which one OSN responded almost exclusively to o-cresol, with minor responses to structurally related compounds such as acetophenone and guaiacol (Fig. 4c).

#### d. Loss of response

In *Dmel*, the Mab4A OSN responds to E2-hexanal, a green leaf volatile. This response was absent in both *Dvir* and *Dbus* (red arrowheads, Fig. 4d). In contrast, the neighboring ab4B OSN, which encodes geosmin, was strongly conserved across all three species.

## Discussion

In this study, I aimed to compare the antennal basiconic olfactory coding across three drosophilid species representing three subgenera and occupying ecologically distinct niches. I found that the coding of the same set of odorants by the antennal basiconic sensilla is partially conserved, with modest similarities between *Dmel* and *Dbus* but no detectable similarity when compared with *Dvir*. This finding is notable given the distinct natural habitats of the three species [23–25,29]. Although their ecological habitats show highly diverse and non-overlapping chemical profiles (Fig. 1b), peripheral basiconic coding exhibited partial conservation across *Dmel* and *Dbus*. This observation is broadly consistent with previous studies showing that some features of peripheral global coding remain conserved despite ecological divergence in drosophilids [17,19,30]. This study extends these findings by including phylogenetically more distant species from distinct ecological niches.

I restricted the scope of the analysis to the antennal basiconic sensilla, consistent with previous studies, as they represent the most chemically diverse class of olfactory sensilla and are primarily involved in host odor detection. Because sensilla of the maxillary palp also contribute to host odor detection [31] and were not included in this study, the conclusions presented here reflect antennal basiconic coding patterns rather than the global olfactory coding landscape.

Importantly, prior work has demonstrated that changes in individual circuits, whether at the receptor or circuit level, can drive discrimination of niche-defining cues and host specialization [2,6,12,16,32]. Consistent with this, ligands such as dimethyldisulfide (DMDS) [7] and guaiacol [8] were identified as key host-signifying cues in *Dbus* and *Dvir*, respectively, that failed to induce significant behavior in *Dmel* and were sufficient to elicit host-specialising behaviors. Together, these results suggest that evolutionary shifts at the level of individual circuits likely underlie host specialization.

I also acknowledge that I have selected a single species as a representative of each subgenus; therefore, care must be taken when extrapolating these findings to general evolutionary trends across subgenera. Another important limitation of this study is that, although I match OSN types based on their functional properties, the receptor identities and true orthology remain unresolved. Future work involving receptor identification and circuit mapping (in *Dvir* and *Dbus*) will be needed to determine whether the observed conserved response profiles result from homologous OSN types or convergent response evolution.

I identified several compounds that are coded in a highly conserved manner across the peripheral olfactory systems of the three species investigated (Fig. 2). In particular, OSNs detecting key ligands such as guaiacol and geosmin showed strong conservation at the peripheral level. However, despite this conservation, the three species exhibited divergent behavioral responses to the same ligands. This suggests that factors beyond peripheral coding, likely involving higher-order processing and non-neuronal cells, play a critical role in shaping the final behavioral outcome [33,34]. A similar conclusion was drawn in a recent study showing that behavioral responses to identical compounds differed significantly across five ecologically distant drosophilid species [30], pointing toward evolutionary divergence at the circuit level in the brain. I found that only seven out of forty-three tested compounds were detected in a conserved manner, whereas the majority were processed in distinct ways. This suggests that the tuning properties of OSNs may have evolved differently across the three species, for reasons that remain to be elucidated.

Interestingly, many compounds that I tested were behaviorally neutral, even though previous studies have reported clear valences. For instance, guaiacol and methyl salicylate were reported as aversive and attractive, respectively, in *Dvir* [35]. Similarly, methyl salicylate is strongly aversive in *Dmel* [36] but attractive in *Dbus* [37]. A key difference lies in the concentrations tested, as earlier studies used 10^−2^ v/v, whereas I applied a 100-fold lower concentration (10^−4^ v/v), reflecting levels more comparable to those occurring naturally.

I found geosmin to be aversive in both *Dvir* and *Dmel*. Geosmin detection is broadly conserved across drosophilids and serves as an aversive cue in *Dmel*, where it signals the presence of harmful microbes [27]. Surprisingly, *Dbus* did not show aversion to geosmin (Fig. 3a). Instead, the response trended toward attraction, though not significantly. This is noteworthy given that *Dbus* tolerates toxic DMDS and preferentially breeds on DMDS-emitting substrates [7]. Moreover, the species is known to colonize oranges at stages much later than the fermentation stage favored by most drosophilids [38,39]. These observations raise the possibility that *Dbus* could have evolved a broader tolerance to microbe-rich substrates, potentially including those emitting geosmin, and that its shifted niche preference may have facilitated a reduced or altered aversive response to this compound.

I found that in *Dbus*, an OSN with a large action potential amplitude responds to CO_2_, whereas in *Dmel* CO_2_ detection is canonically mediated by a smaller-amplitude OSN [40,41], which is linked to aversion in several drosophilids [42,43]. Previous work has suggested that OSN spike amplitude can influence odor valence, with larger-amplitude OSNs typically mediating attraction and smaller-amplitude OSNs mediating aversion [44]. Notably, CO_2_ is detected by large-spiking cpA OSNs in the maxillary palps of mosquitoes, but by small-spiking Mab1C OSNs in the antennae of *Dmel* [40,41,45]. Thus, detection of CO_2_ by a large-spiking OSN in *Dbus* raises the intriguing possibility that CO_2_ may be attractive in this species, providing an opportunity to study the evolutionary rewiring of the CO_2_ circuit. Alternatively, the difference may reflect anatomical changes at the level of OSN dendrites [46] or mitochondrial density near the inner dendrites rather than alterations in central circuitry.

I found that Vab3A OSNs in *Dvir* respond more strongly to methyl hexanoate than to ethyl hexanoate (Fig. 4b). This mirrors a key shift previously reported in *D. sechellia*, where altered sensitivity contributes to host specialization [2,12]. It is puzzling to observe a similar trend in *Dvir*, given that these esters were not detected in the natural headspace of slime fluxes, its known niche [8]. However, both esters are components of the aggregation pheromones produced by *Dvir* males [47], suggesting that the sensitivity shift may influence intraspecific aggregation rather than niche adaptation.

In *Dbus*, I identified two OSN types responding to o-cresol, despite the compound being behaviorally neutral. In nature, o-cresol is found in beaver castoreum and is a strong attractant in *Dvir* [8,23], but it was absent from the headspaces of rotting vegetables, the primary niche of *Dbus*. The additional rare OSN type (Bab10) responded to both acetophenone and o-cresol, raising the possibility that a structurally similar phenolic present in vegetable headspaces may be its true ecological ligand. Such redundancy may improve discrimination between closely related odorants. A comparable case was recently described in *Dmel*, where a newly defined OSN type (ab11A) responded to both o-cresol and indole, while the canonical ab6B OSN responded to o-cresol but not to indole [48].

I found that Vab4A and Bab4A OSNs do not respond to E2-hexanal. E2-hexanal in *Dmel* is detected by Or7a [28]. However, this odorant receptor is not expressed in *Dvir* [11], supporting the observed absence of response, while no information about the same is available in the case of *Dbus*. Furthermore, several diagnostic ligands identified in our screening elicited only weak responses (less than 25 Hz), indicating that the optimal ligands for these OSN types and their behavioral contributions have yet to be identified and remain an important direction for future studies.

Taken together, an important implication of this study is that conservation at the peripheral level does not necessarily predict conservation at the behavioral level. Even when the overall strengths of functionally matched OSNs do not differ, the behavioral valence of the same ligands can vary among species. These results suggest that host specialization is driven less by shifts in the global olfactory code and more by evolutionary changes within individual circuits and higher-order processing beyond peripheral detection.

## Materials and methods

### *Drosophila* stock

Wild-type species [*D. busckii*: 13000-0081.00, *D. virilis*: 15010-1051.00 and *D. melanogaster* CantonS] were used in this study. *D. busckii* were maintained on Wheeler-Clyton (Double-layer food; 2:0.5:0.2 ratio) while *D. virilis* and *D. melanogaster* were kept on standard fly food ([7], supplementary table). All three species were maintained at 12:12 h light: dark cycle at 23⍰C and at 40% relative humidity and were in the lab for more than 5 years at the time of testing.

### Chemical stimuli

All chemicals used in this study were purchased with the highest purity possible. A list of all odorants used, along with their suppliers, is available in supplementary information [7]. Dilutions were done in mineral oil while testing in behavioral bioassays, and were diluted to a final concentration of 10^-4^ v/v.

### Behavioral bioassays

Two choice assays were conducted as previously described [8]. In brief, flies of both sexes were kept together for 6 days post-eclosion. A mixed group of 7–9-day-old flies was used for behavioral studies. A group of 35 flies was used for experiments conducted in salad boxes (transparent plastic boxes, ∼5*∼7*∼10 cm (w*l*h) with 10 ventilation holes punctured with forceps). Flies were sorted one day before the experiment using CO_2_ pads and starved on 500 μl distilled water *ad libitum*. Experiments generally began around 1100 hrs. and were terminated around the same time after 48 hours with a 16L: 8D photoperiod during testing. The preference index was calculated as (T-C)/(T+C), where T represents the number of flies in the test trap while C represents the same in the control trap. Traps were manually created by attaching pink paper cones to plastic vials.

### Electrophysiology

Single sensillum recording (SSR) data used in this study was obtained from experiments carried out in two publicly available studies [7,8]. SSR were performed by following a protocol previously described in detail elsewhere [49]. Typically, 5 to 10-day-old female flies were used in the experiment. All odorants for the antennal screening experiment were diluted in hexane and tested at a concentration of 10^-4^ (v/v) unless stated otherwise. For dose-response curve experiments, a freshly prepared serial dilution of both esters was carried out. CO_2_ was delivered as a manual exhalation by the researcher. Diluted odorants were pipetted into an odor cartridge described previously [49], and the same cartridge was used not more than 3 times for dose-response and antenna screening experiments, respectively, unless stated otherwise. For all recordings, a spike change of more than 25 Hz was defined as a strong response and used as a diagnostic stimulus to identify the corresponding OSN class.

### Statistical analysis

First, odor to odor correlation matrices were generated between each species pair using the raw SSR response data generated across all sensillum types. Spearman correlations were calculated between all pairs of odors across all OSN types. Because absolute firing rates carry biologically relevant information, analyses were performed on raw responses, and no per-OSN normalization or batch effect correction was applied before clustering. Similarity between species in the global odor coding was further compared by using a Mantel test with Spearman correlation and 10,000 permutations. To estimate uncertainty, odors were resampled using a bootstrap value of 10000 and 95% confidence levels were calculated from the bootstrap distributions. Finally, p-values were corrected across the three species pair comparisons using the Holm method. For hierarchical clustering analysis, first a distance matrix was calculated using Euclidean distance followed by clustering using the Ward.D2 method in R. Statistical analyses pertaining to Figure 3 were performed using GraphPad-Prism 9.1.1 (https://www.graphpad.com/scientific-software/prism/). SSR traces were analysed using AutoSpike32 software version 3.7 (Syntech, NL 1998). Changes in action potential (spike count) were calculated by subtracting the number of spikes one second before (spontaneous activity) from those elicited one second after the onset of the stimulus. For behavioural data analyses, the raw data count was converted to an index. Such index replicates were first tested for normal (Gaussian) distribution using the Shapiro-Wilk normality test (significant = 0.05). Most of the data was observed to be normally distributed. Preference indices were tested against neutrality using one-sample t-tests when normally distributed, or non-parametric alternatives otherwise. Where comparisons were made between species or groups, Welch’s t-test was used. Graphs were generated using GraphPad Prism 9.1.1. and figures were constructed and processed with Adobe Illustrator CS5 and Adobe Photoshop (Adobe system Inc.). A principal component analysis of all chromatograms was generated by using an online software called XCMS version 3.7.1 [26].

## Supporting information

Diagnostic odors

## Ethics

This work did not require ethical approval from a human subject or animal welfare committee.

## Acknowledgements

I am grateful to Prof. Dr Bill S. Hansson for providing access to the departmental resources at the Max Planck Institute for Chemical Ecology, for his supervision during the work on this manuscript, and for comments on the first draft of the manuscript. I thank Dr Markus Knaden for his support. I thank Silke Trautheim, Roland Spiess, Ibrahim Alali and Manal Alali for their help with maintaining fly stocks. I also thank Swetlana Laubrich for administrative assistance.

## Declaration of AI use

I acknowledge the assistance of ChatGPT (OpenAI) for suggesting edits aimed at improving the grammar and clarity of the first version of the text. Pre-submission review was conducted using QED science (https://www.qedscience.com/)

## Authors’ contribution

V.P.M. conceived the project, designed the experiments, conducted all experiments, made the figures, analyzed data, and wrote the manuscript.

## References

1. Pal Mahadevan V: Exploring the olfactory neuroecology of non-model drosophilids. 2024,

2. Dekker T, Ibba I, Siju KP, Stensmyr MC, Hansson BS: Olfactory shifts parallel superspecialism for toxic fruit in Drosophila melanogaster sibling, D. sechellia. Current Biology 2006, 16:101–109.

3. Stensmyr MC, Giordano E, Balloi A, Angioy AM, Hansson BS: Novel natural ligands for Drosophila olfactory receptor neurones. Journal of Experimental Biology 2003, 206:715– 724.

4. Keesey IW, Knaden M, Hansson BS: Olfactory Specialization in Drosophila suzukii Supports an Ecological Shift in Host Preference from Rotten to Fresh Fruit. J Chem Ecol 2015, 41:121–128.

5. Crowley-Gall A, Date P, Han C, Rhodes N, Andolfatto P, Layne JE, Rollmann SM: Population differences in olfaction accompany host shift in Drosophila mojavensis. Proc Biol Sci 2016, 283.

6. Linz J, Baschwitz A, Strutz A, Dweck HKM, Sachse S, Hansson BS, Stensmyr MC: Host plant-driven sensory specialization in Drosophila erecta. Proceedings of the Royal Society B: Biological Sciences 2013, 280.

7. Pal Mahadevan V, Galagovsky D, Knaden M, Hansson BS: Preference for and resistance to a toxic sulfur volatile opens up a unique niche in Drosophila busckii. Nature Communications 2025 16:1 2025, 16:1–14.

8. Mahadevan VP, Stieber-Rödiger R, Knaden M, Hansson BS: Phenolics as ecologically relevant cues for slime flux breeding Drosophila virilis. iScience 2024, 27:111180.

9. Gardiner A, Barker D, Butlin RK, Jordan WC, Ritchie MG: Drosophila chemoreceptor gene evolution: selection, specialization and genome size. Mol Ecol 2008, 17:1648–1657.

10. McBride CS, Arguello JR: Five Drosophila genomes reveal nonneutral evolution and the signature of host specialization in the chemoreceptor superfamily. Genetics 2007, 177:1395–1416.

11. Guo S, Kim J: Molecular Evolution of Drosophila Odorant Receptor Genes. Mol Biol Evol 2007, 24:1198–1207.

12. Auer TO, Khallaf MA, Silbering AF, Zappia G, Ellis K, Álvarez-Ocaña R, Arguello JR, Hansson BS, Jefferis GSXE, Caron SJC, et al.: Olfactory receptor and circuit evolution promote host specialization. Nature 2020, 579:402–408.

13. Ramasamy S, Ometto L, Crava CM, Revadi S, Kaur R, Horner DS, Pisani D, Dekker T, Anfora G, Rota-Stabelli O: The Evolution of Olfactory Gene Families in Drosophila and the Genomic Basis of chemical-Ecological Adaptation in Drosophila suzukii. Genome Biol Evol 2016, 8:2297–2311.

14. Keesey IW, Zhang J, Depetris-Chauvin A, Obiero GF, Gupta A, Gupta N, Vogel H, Knaden M, Hansson BS: Functional olfactory evolution in Drosophila suzukii and the subgenus Sophophora. iScience 2022, 25:104212.

15. Xue Q, Dweck HKM: Receptor sequence divergence, gain, loss, duplication, and neofunctionalization drive olfactory adaptation in Drosophila suzukii. Proc Natl Acad Sci U S A 2026, 123:e2529586123.

16. Zhao Z, McBride CS: Evolution of olfactory circuits in insects. J Comp Physiol A Neuroethol Sens Neural Behav Physiol 2020, 206:353–367.

17. De Bruyne M, Smart R, Zammit E, Warr CG: Functional and molecular evolution of olfactory neurons and receptors for aliphatic esters across the Drosophila genus. J Comp Physiol A 2010, 196:97–109.

18. Depetris-Chauvin A, Galagovsky D, Keesey IW, Hansson BS, Sachse S, Knaden M: Evolution at multiple processing levels underlies odor-guided behavior in the genus Drosophila. Current Biology 2023, 33:4771-4785.e7.

19. Stensmyr MC, Dekker T, Hansson BS: Evolution of the olfactory code in the Drosophila melanogaster subgroup. Proc R Soc Lond B Biol Sci 2003, 270:2333–2340.

20. Shaw KH, Johnson TK, Anderson A, De Bruyne M, Warr CG: Molecular and Functional Evolution at the Odorant Receptor Or22 Locus in Drosophila melanogaster. Mol Biol Evol 2019, 36:919–929.

21. Keesey IW, Zhang J, Depetris-Chauvin A, Obiero GF, Gupta A, Gupta N, Vogel H, Knaden M, Hansson BS: Functional olfactory evolution in Drosophila suzukii and the subgenus Sophophora. iScience 2022, 25:104212.

22. Takagi S, Sancer G, Abuin L, Stupski SD, Roman Arguello J, Prieto-Godino LL, Stern DL, Cruchet S, Álvarez-Ocaña R, Wienecke CFR, et al.: Olfactory sensory neuron population expansions influence projection neuron adaptation and enhance odour tracking. Nature Communications 2024 15:1 2024, 15:1–18.

23. Spieth HT: The Virilis Group of Drosophila and the Beaver Castor. Source: The American Naturalist 1979, 114:312–316.

24. Hoikkala A, Poikela N: Adaptation and ecological speciation in seasonally varying environments at high latitudes: Drosophila virilis group. Fly (Austin) 2022, 16:85–104.

25. Atkinson W: Ecological studies of the breeding sites and reproductive strategies of domestic species of Drosophila. 1977.

26. Tautenhahn R, Patti GJ, Rinehart D, Siuzdak G: XCMS Online: A Web-Based Platform to Process Untargeted Metabolomic Data. 2012, doi:10.1021/ac300698c.

27. Stensmyr MC, Dweck HKM, Farhan A, Ibba I, Strutz A, Mukunda L, Linz J, Grabe V, Steck K, Lavista-Llanos S, et al.: A Conserved Dedicated Olfactory Circuit for Detecting Harmful Microbes in Drosophila. Cell 2012, 151:1345–1357.

28. Hallem EA, Carlson JR: Coding of Odors by a Receptor Repertoire. Cell 2006, 125:143–160.

29. Mansourian S, Stensmyr MC: The chemical ecology of the fly. Curr Opin Neurobiol 2015, 34:95–102.

30. Depetris-Chauvin A, Galagovsky D, Keesey IW, Hansson BS, Sachse S, Knaden M: Evolution at multiple processing levels underlies odor-guided behavior in the genus Drosophila. Current Biology 2023, doi:10.1016/J.CUB.2023.09.039.

31. De Bruyne M, Clyne PJ, Carlson JR: Odor coding in a model olfactory organ: the Drosophila maxillary palp. J Neurosci 1999, 19:4520–4532.

32. Prieto-Godino LL, Rytz R, Cruchet S, Bargeton B, Abuin L, Silbering AF, Ruta V, Dal Peraro M, Benton R: Evolution of Acid-Sensing Olfactory Circuits in Drosophilids. Neuron 2017, 93:661-676.e6.

33. Seki Y, Dweck HKM, Rybak J, Wicher D, Sachse S, Hansson BS: Olfactory coding from the periphery to higher brain centers in the Drosophila brain. BMC Biol 2017, 15:1–20.

34. Lee D, Shahandeh MP, Abuin L, Benton R: Comparative single-cell transcriptomic atlases of drosophilid brains suggest glial evolution during ecological adaptation. PLoS Biol 2025, 23.

35. Baleba SBS, Mahadevan VP, Knaden M, Hansson BS: Temperature-dependent modulation of odor-dependent behavior in three drosophilid fly species of differing thermal preference. Communications Biology 2023 6:1 2023, 6:1–11.

36. Knaden M, Strutz A, Ahsan J, Sachse S, Hansson BS: Spatial Representation of Odorant Valence in an Insect Brain. Cell Rep 2012, 1:392–399.

37. Buda V, Radžiute S, Lutovinovas E: Attractant for Vinegar Fly, Drosophila busckii, and Cluster Fly, Pollenia rudis (Diptera: Drosophilidae et Calliphoridae). Zeitschrift fur Naturforschung - Section C Journal of Biosciences 2009, 64:267–270.

38. Nunney L: Drosophila on oranges: colonization, competition, and coexistence. Ecology 1990, 71:1904–1915.

39. Nunney L, Nunney L: The colonization of organges by the cosmopolitan Drosophila. Oecologia 1996 108:3 1996, 108:552–561.

40. Jones WD, Cayirlioglu P, Grunwald Kadow I, Vosshall LB: Two chemosensory receptors together mediate carbon dioxide detection in Drosophila. Nature 2007, 445:86–90.

41. Kwon JY, Dahanukar A, Weiss LA, Carlson JR: The molecular basis of CO2 reception in Drosophila. Proc Natl Acad Sci U S A 2007, 104:3574–3578.

42. van Loon JJA, Smallegange RC, Bukovinszkiné-Kiss G, Jacobs F, De Rijk M, Mukabana WR, Verhulst NO, Menger DJ, Takken W: Mosquito Attraction: Crucial Role of Carbon Dioxide in Formulation of a Five-Component Blend of Human-Derived Volatiles. J Chem Ecol 2015, 41:567.

43. Pan JW, McLaughlin J, Yang H, Leo C, Rambarat P, Okuwa S, Monroy-Eklund A, Clark S, Jones CD, Volkan PC: Comparative analysis of behavioral and transcriptional variation underlying CO2 sensory neuron function and development in Drosophila. Fly (Austin) 2017, 11:239–252.

44. Wu ST, Chen JY, Martin V, Ng R, Zhang Y, Grover D, Greenspan RJ, Aljadeff J, Su CY: Valence opponency in peripheral olfactory processing. Proc Natl Acad Sci U S A 2022, 119:e2120134119.

45. Tauxe GM, Macwilliam D, Boyle SM, Guda T, Ray A: Targeting a dual detector of skin and CO2 to modify mosquito host seeking. Cell 2013, 155:1365.

46. Gonzales CN, McKaughan Q, Bushong EA, Cauwenberghs K, Ng R, Madany M, Ellisman MH, Su CY: Systematic morphological and morphometric analysis of identified olfactory receptor neurons in drosophila melanogaster. Elife 2021, 10.

47. Bartelt RJ, Jackson LL, Schaner AM: Ester components of aggregation pheromone of Drosophila virilis (Diptera: Drosophilidae). J Chem Ecol 1985, 11:1197–1208.

48. Benton R, Mermet J, Jang A, Endo K, Cruchet S, Menuz K: An integrated anatomical, functional and evolutionary view of the Drosophila olfactory system. EMBO Rep 2025, 26:3204–3225.

49. Mahadevan VP, Lavista-Llanos S, Knaden M, Hansson BS: No functional contribution of the gustatory receptor, Gr64b, co-expressed in olfactory sensory neurons of Drosophila melanogaster. Front Ecol Evol 2022, 10:869.

